# The serine/threonine kinase CcAkt regulates the fertility of *Coridius chinensis*

**DOI:** 10.1101/2023.10.07.561341

**Authors:** Jinyu Feng, Juan Du, Shangwei Li, Xingxing Chen

## Abstract

Akt (also known as protein kinase B) belongs to the multifunctional serine/threonine kinase family and is an important component of the insulin signaling pathway that plays a key role in many biological processes such as cell growth, proliferation, and survival. However, few studies have reported the effect of Akt on reproduction in Hemiptera. In this study, we cloned and characterized the *Akt* gene from *Coridius chinensis* (*CcAkt*). The open reading frame of *CcAkt* has a length of 1,563 bp and encodes 520 amino acids. It has a conserved pleckstrin homology domain, a serine/threonine protein kinase catalytic domain, and a serine/threonine protein kinase domain. Phylogenetic analysis showed that *CcAkt* and *HhAkt* of *Halyomorpha halys* had the highest similarity. Analysis of temporal and spatial expression patterns revealed that *CcAkt* is expressed throughout development and in various tissues of *C. chinensis* adults. *CcAkt* expression was high in female adult and the fourth instar nymph stage of *C. chinensis*. In *C. chinensis* adult, *CcAkt* expression was highest in the testis and ovary. Injection of bovine insulin induced *CcAkt* expression, whereas that of 20-hydroxyecdysone significantly reduced *CcAkt* expression. Both hormones, however, induced the expression of *vitellogenin* (*Vg*) and *vitellogenin receptor* (*VgR*). In unmated females, *CcAkt* knockout resulted in decreased expression of *CcVg* and *CcVgR*, stunted the development of the ovarioles, decreased the number of eggs and hatching rate. These findings suggest that CcAkt may be involved in regulating the reproduction of *C. chinensis*.

## 1. Introduction

The insulin signaling pathway, a conserved signaling pathway, is present in common model organisms such as *Drosophila melanogaster* and *Bombyx mori*, and it is structurally and functionally conserved [1,2]. The insulin signaling pathway has different roles in different developmental stages of insects; there are numerous studies showing that insulin can regulate insect cell proliferation, differentiation and development, as well as insect reproduction and lifespan through two signaling pathways, the Ras–MAPK pathway and the PI3K–Akt pathway [3-5]. Akt (also known as protein kinase B) belongs to a family of multifunctional serine/threonine kinases that act as central molecules in the insulin signaling pathway, binding and phosphorylating downstream proteins and transcription factors, thereby altering their subcellular localization and controlling their activation or inhibition to participate in the regulation of various physiological activities in insects [6,7]. In *D. melanogaster*, the insulin signaling pathway directly regulates oocyte growth, and silencing key factors in the insulin signaling pathway prevents the uptake of yolkogenic protein precursors, ultimately leading to retarded oocyte development [8]. In *Maruca vitrata*, silencing of *MvAkt* suppressed the *Vg* (*vitellogenin*) and *VgR* (*vitellogenin receptor*) expression and prevented normal ovarian development [9]. In *Chrysopa pallens*, the knock-down of *CpAkt* by RNA interference (RNAi) significantly suppressed *Vg* expression and led to ovarian atrophy and reduced yolk protein deposition [10]. *BmAkt* was also associated with the embryonic lagging process in *B. mori* [11].

Previous studies have found that endocrine hormones are involved in and regulate many physiological and developmental processes in insects [12-14]. The endocrinology of insect reproduction has been studied with respect to three hormones, insulin, juvenile hormones (JH), and ecdysone; these three hormones are also closely related to the insect insulin signaling pathway [15,16]. JH is synthesized and secreted by the corpus and is involved in the regulation of insect metamorphosis and reproduction [17,18]. Similarly, insulin-like peptides (ILPs), encoded by a multigene family expressed in the brain and other tissues can also regulate numerous physiological processes in insects [2]. In insects, ILPs regulate nutritional status, insect growth, development, longevity, reproduction, and other physiological processes through a conserved insulin signaling pathway [19-23]. It has been shown that the injection of bovine insulin into *Tribolium castaneum* female adults significantly increased the *Vg* mRNA and protein expression, suggesting that ILPs are involved in regulating *Vg* synthesis [17]. In the prothorax of *B. mori*, bovine insulin enhances the phosphorylation of *BmAkt* and stimulates the secretion of ecdysone [24]. Ecdysone, a typical steroid hormone, is produced in the prothoracic gland of the larval stage of insects, but the prothoracic gland is degraded in the adult stage, so juvenile hormone and ecdysone are synthesized in the gonads, and these two hormones promote gonad maturation and play a role in the regulation of insect reproduction [25,26]. In *Aedes aegypti*, bovine insulin stimulated the ovarian secretion of ecdysone, and a bovine insulin concentration of 17 μM was optimal to induce this effect [27]. *In vitro* incubation of *A. aegypti* isolated adiposomes with insulin and ecdysone activates *Vg* gene expression, and downregulation of *AaAkt* inhibits *Vg* activation through insulin and ecdysone [28]. These findings suggest that the insulin signaling pathway and the three exogenous hormones are mutually regulated to influence the growth and developmental processes of insects.

*Coridius chinensis* belongs to the family Dinidoridae of the order Hemiptera and is a traditional resource insect with great developmental value. The resource of wild *C. chinensis* is in short supply; moreover, research into artificial breeding technology is not sufficiently well developed and there are many challenges facing large-scale breeding. Therefore, this study was conducted to investigate the effects of insulin signaling pathway and exogenous hormones on the reproductive ability of *C. chinensis*, to provide a theoretical basis for the artificial large-scale breeding of *C. chinensis*. In this study, we cloned and characterized the open reading frame (ORF) of *Akt* from *C. chinensis* (*CcAkt*). The spatio-temporal expression pattern of *CcAkt* was analyzed by using real-time quantitative PCR (RT-qPCR); the role of *CcAkt* in the reproductive regulation of *C. chinensis* was investigated using RNAi technology; and the effects of injections of bovine insulin and ecdysone on *CcAkt* and reproduction-related genes were explored.

## 2. Materials and Methods

### 2.1. Insects

*C. chinensis* was harvested from rice fields in Guiyang city, Guizhou province, China, and reared in an artificial climate chamber with fresh pumpkin leaves. The temperature of the artificial climate chamber was 28°C ±1°C, the relative humidity was 80%±5%, and the photoperiod was 16 h light: 8 h dark (16L: 8D).

### 2.2. Gene Cloning

Total RNA of *C. chinensis* was extracted using an Easter Super Total RNA Extraction Kit (Promega, Madison, WI, USA). cDNA was synthesized from the above total extracted RNA as a template using a PrimeScript First-Strand cDNA Synthesis Kit (Takara Bio, Beijing, China). Specific primers (Table 1) were designed using Primer Premier 6.0 (PREMIER Biosoft, Palo Alto, CA, USA) based on the sequence information for the *C. chinensis* transcriptome. Reverse transcription-polymerase chain reaction (RT-PCR) was used to amplify the target fragment from the synthesized cDNA as a template. The amplification products were then purified using a DiaSpin DNA Gel Extraction Kit (Sangon Biotech, Shanghai, China), recovered, and cloned into pMD19-T vector (Takara Bio, Dalian, China) before transformation into *E. coli* DH5α competent cells (Invitrogen, Carlsbad, CA, USA) in overnight culture. Single colonies were picked from the overnight culture plates for PCR amplification confirmation and the correct recombinant plasmids were sent to Sangon Biotech (Shanghai, China) for sequencing.

**Table 1.**
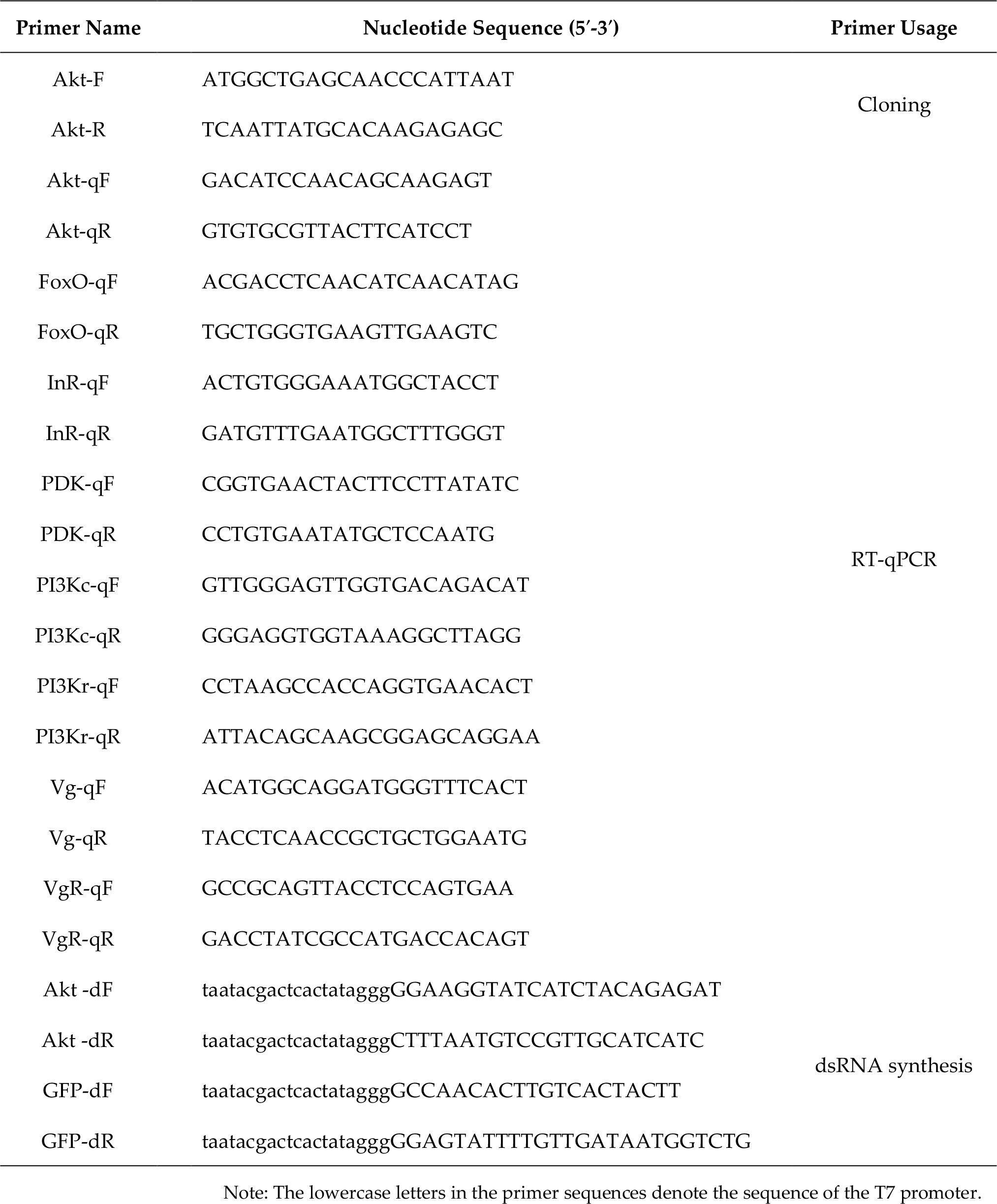
Primers used for cloning, expression analysis and RNA interference of CcAkt from *C. chinensis*.

### 2.3. Bioinformatic Analyses of CcAkt

The ORF of *CcAkt* was confirmed with ORFfinder (https://www.ncbi.nlm.nih.gov/orffinder). The molecular weight and isoelectric point of CcAkt were predicted by using the Expasy ProtParam platform (https://web.expasy.org/protparam). Domains of this protein were analyzed with InterPro (https://www.ebi.ac.uk/interpro). Prediction of glycosylation sites was carried out by employing NetOGlyc-4.0 (https://services.healthtech.dtu.dk/service.php?NetOGlyc-4.0) and NetNGlyc-1.0 (https://services.healthtech.dtu.dk/service.php?NetNGlyc-1.0). Three-dimensional (3D) structure of CcAkt was constructed with SWISS-MODEL homologous modeling (https://swissmodel.expasy.org) and PyMOL 2.5 (Schrodinger, New York, NY, USA) was used to draw its molecular structure. A phylogenetic tree was constructed by using the neighbor-joining method in the MEGA 11 software with 1,000 runs.

### 2.4. Gene Expression Analyses

Samples were collected from eight different developmental stages of *C. chinensis* (eggs, first to fifth instar nymphs, female adults, and male adults) and seven different tissues (the head, integument, fat body, hemolymph, midgut, testis, ovary, and muscle); three biological replicates were used. Total RNA extraction and cDNA synthesis were performed as described above. RT-qPCR primers (Table 1) were designed using Premier 6.0 (PREMIER Biosoft, Palo Alto, CA, USA), and *CcAkt* mRNA expression was normalized using *β-actin* (GenBank ID: MK370101) as the internal reference gene. The reaction system comprised 1-μL cDNA, 1 μL each of forward and reverse primers, 10-μL TB Green Premix Ex Taq II (Takara, Beijing, China), and 7-μL sterile deionized water. The reaction parameters were as follows: 50°C for 2 min; 95°C for 2 min; followed by 40 cycles of 95°C for 15 s and 60°C for 1 min.

### 2.5. Hormone Treatment

Bovine insulin powder (0.2 g, purity: ≥ 27 USP units/mg) (Solarbio, Beijing, China) was weighed and dissolved in 20 mL of 25 mM HEPES buffer. Then, 10 mg/mL of bovine insulin mother liquor was obtained by filtration and sterilization. The prepared bovine insulin master mix was diluted to 2,000 ng/μL and 1 μL of this mix per insect was injected to the body cavity via the metapleuron under the wing using a Nanoliter (WPI, Sarasota, FL, USA). An equal amount of HEPES buffer was injected as a control to negate the effects caused by mechanical damage. For the 20E treatment, 1 mg of 20E (HPLC: ≥ 95% purity) (Solarbio, Beijing, China) powder was dissolved in 15 μL of 95% ethanol to obtain a final concentration of 100 μg/μL of 20E master mix. The master mix is diluted to 2,000 ng/μl and 1 μl of this mix is injected per insect into the female adult.; an equal amount of 0.1% anhydrous ethanol was injected as the control. Three biological replicates were performed, with 20 animals per group. The injected female adults were placed in an artificial climate chamber (a temperature of 28°C ± 1°C, relative humidity of 80% ± 5%, and a 16L: 8D photoperiod). Three female adults from each of the treatment and control groups were collected at 3, 6, 12 and 24 h after injection. *CcAkt, CcVg*, and *CcVgR* expressions were detected by using RT-qPCR.

### 2.6. RNA Interference

Primers (Table 1) for RNAi were designed according to siDirect (http://sidi-rect2.rnai.jp) and DSIR (http://biodev.extra.cea.fr/DSIR/DSIR.html), and ds*CcAkt* was prepared and purified using a TranscriptAid T7 High Yield Transcription Kit (Thermo Fisher Scientific, Waltham, MA, USA). The purified dsRNA was diluted to 2,000 ng/μL and 3 μg of dsRNA were injected with a Nanoliter (WPI, Sarasota, FL, USA) to the body cavity via the metapleuron under the wing. Each insect was injected with the same amount of *green fluorescent protein* (*GFP*) dsRNA from *Aequorea victoria* (GenBank accession number: CAA58789) as a negative control. Each group included 20 insects and three biological replicates were performed. The dsRNA-injected female adults were reared in an artificial climate chamber under the normal conditions. At 24, 48, 72 and 96 h after injection, female individuals were collected and the efficiency of RNAi was detected with RT-qPCR. The group with the highest decrease in *CcAkt* expression level among the above four experimental groups was taken for the follow-up experiment. Analysis of mRNA expression levels of five key genes related to insulin signaling pathway and reproductive regulators *CcVg* and *CcVgR* after ds*CcAkt* injection. To determine the effect of *CcAkt* on the reproductive capacity of *C. chinensis*, we dissected and photographed the ovaries using a VHX-2000 super-resolution digital microscope (Keyence, Tokyo, Japan) and measured the length of the ovarioles in three biological replicates on day 15 after injection of dsRNA. Males were paired with dsRNA-treated virgin females and reared in an artificial climate chamber. Average egg production and hatchability per female were counted 15 days after injection of dsRNA.

### 2.7. Statistical Analyses

Data were presented as the mean ± SD from at least three independent experiments. SPSS 26.0 (SPSS Inc., Chicago, IL, USA) was used for one-way analysis of variance and Duncan’s test was used for multiple comparisons, with *p* < 0.05 indicating a significant difference.

## 3. Results

### 3.1. Characterization of CcAkt

The ORF of *CcAkt* is 1,563 bp in length and encodes 520 amino acids (Figure 1A). ProtParam predicts the relative molecular weight of this nucleotide-derived protein as 59.12 kDa and the isoelectric point as 5.57. The protein contains 75 negatively charged amino acid residues (Asp + Glu) and 63 positively charged amino acid residues (Arg + Lys). The aliphatic amino acid index was 78.48 and the hydrophilic mean coefficient was −0.41, predicting a hydrophilic protein. NetNGlyc and NetOGlyc predicted that the protein has 28 O-glycosylation sites and 2 N-glycosylation sites. Using the NetPhos 3.1 Server to predict the functional sites of the protein online, the protein sequence had 29 possible phosphorylation sites for serine residues, 26 possible phosphorylation sites for threonine residues, and 8 possible phosphorylation sites for tyrosine residues. The amino acid sequence of CcAkt has the conserved structural domains of insect protein kinase B family genes: pleckstrin homology structural domain (PH), serine/threonine protein kinase catalytic structural domain (S_TKc), and serine/threonine protein kinase structural domain (S_TK_C) (Figure 1B). CcAkt forms a monomer consisting of 15 α-helices, 14 β-pleated sheets, and 27 random coils, based on the AlphaFold DB model of Akt1 from *Cryptotermes secundus* (Figure 2A). Evolutionarily conserved amino acids and their positions are showed in the 3D structure (Figure 2B), indicating that they very important for the structure and function of CcAkt.

**Figure 1.**
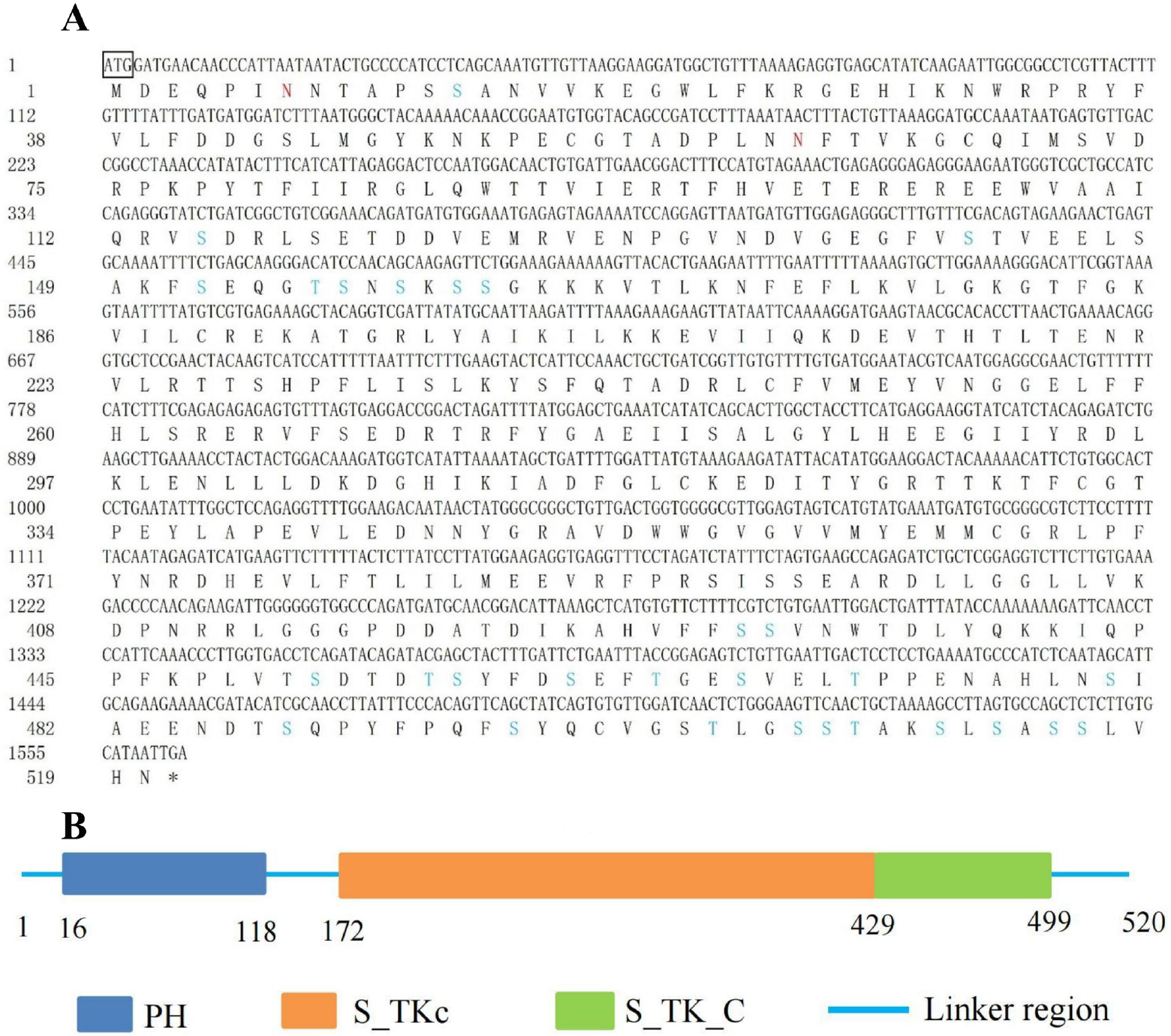
Nucleotide and deduced amino acid sequence and schematic diagram of the domain of *CcAkt* from *C. chinensis*. (A) Nucleotide and amino acid sequences of *CcAkt*. The start codon is indicated with a box and the stop codon is indicated by an asterisk. The N-glycosylation sites are marked with red and the O-glycosylation sites are marked with blue. (B) Schematic diagram of the domain of CcAkt. The blue box, orange box, green box, and blue line represent the pleckstring homology domain (PH), serine/threonine protein kinase catalytic domain (S_TKc), serine/threonine protein kinase domain (S_TK_C), and linker region, respectively.

**Figure 2.**
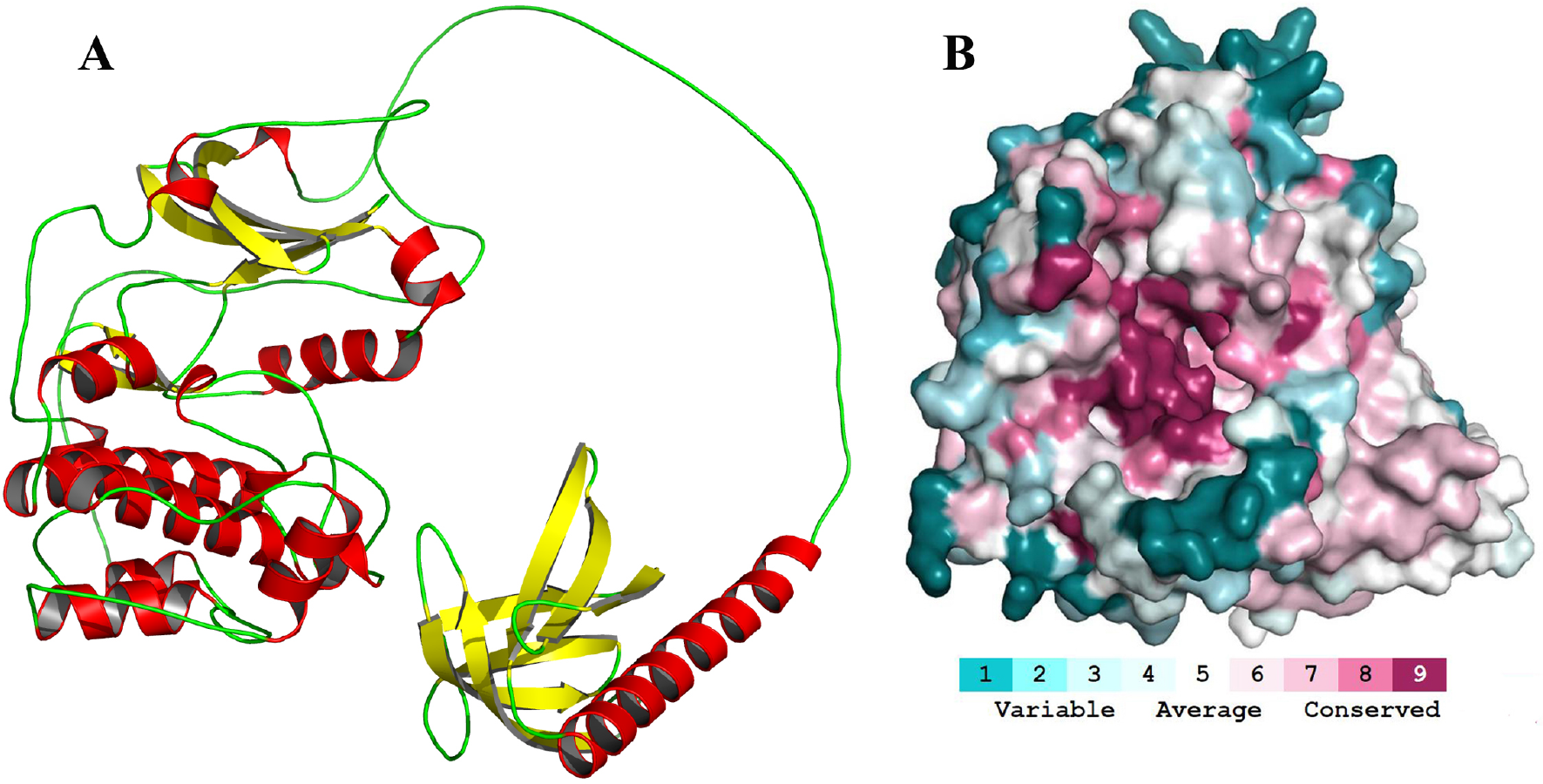
The 3D structure and evolutionary conservation profile of CcAkt. (A) CcAkt forming the molecular structure of a monomer. Red represents α-helices, yellow indicates β-pleated sheets, and green denotes random coils. Homology modeling was performed using AlphaFold v2 based on the AlphaFold DB model of Akt1 from *Cryptotermes secundus*. (B) Highly conserved amino acids and their positions (in deep red) of CcAkt. The ConSurf Server (https://consurf.tau.ac.il) was used to estimate the evolutionary conservation of amino acid positions in CcAkt and 3D graphics were visualized by employing PyMOL 2.5.

### 3.2. Homology Comparison and Cluster Dendrogram

CcAkt was clustered with Akts from 15 other insects and the neighbor-joining method was used to construct a phylogenetic tree. The clustering analysis showed that *C. chinensis* clustered with the *Halyomorpha halys, Cimex lectularius*, and *Homalodisca vitripennis*, indicating that the insect Akt proteins are evolutionarily conserved. *C. chinensis* and *H. halys* were the closest relatives (Figure 3).

**Figure 3.**
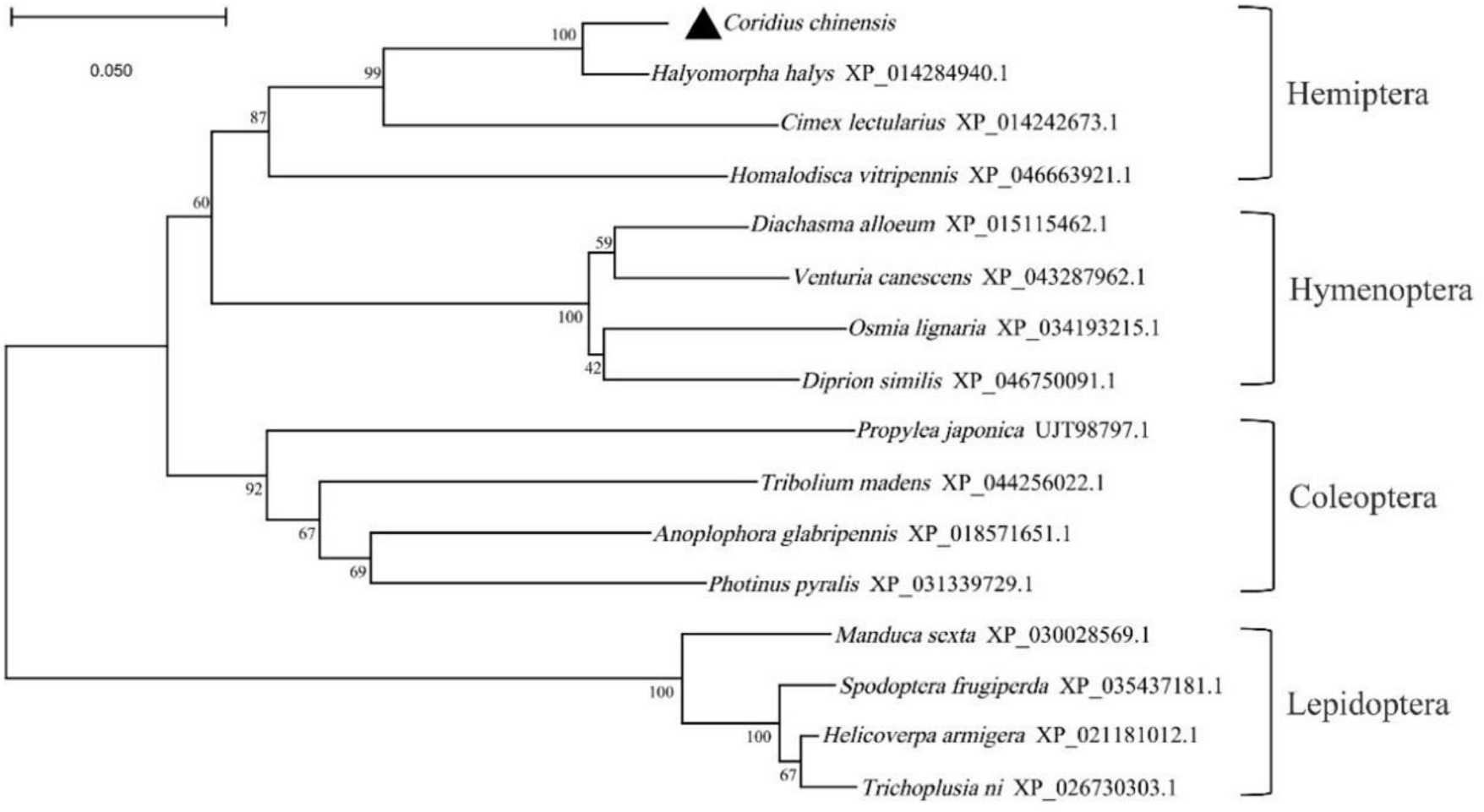
Phylogenetic relationship with Akt of different insects. The phylogenetic tree was generated with MEGA 11 by using neighbor-joining method with 1,000 bootstrap replications, and the number on the node represents the bootstrap test value. CcAkt from *C. chinensis* is marked with a triangle.

### 3.3. Spatiotemporal Expression Patterns

The RT-qPCR results showed that *CcAkt* was expressed specifically throughout the developmental stages of *C. chinensis*; the highest expression was in female adults, followed by the fourth instar worm and egg stage, whereas lower expression was found in other developmental stages and the lowest was found in male adults. *CcAkt* expression was 12.55, 12.09 and 11.97 times higher in adult females, fourth instars, and eggs, respectively, than in males (Figure 4A). *CcAkt* was expressed in different adult tissues; the highest expression was in the testis, followed by the ovary, and lowest expression was in the integument. *CcAkt* expression in the testis and ovary was 322.38-fold and 297.94-fold higher than in the integument, respectively (Figure 4B). The high expression of *CcAkt* in reproductive organs may be related to the fact that reproductive organs are responsible for producing offspring.

**Figure 4.**
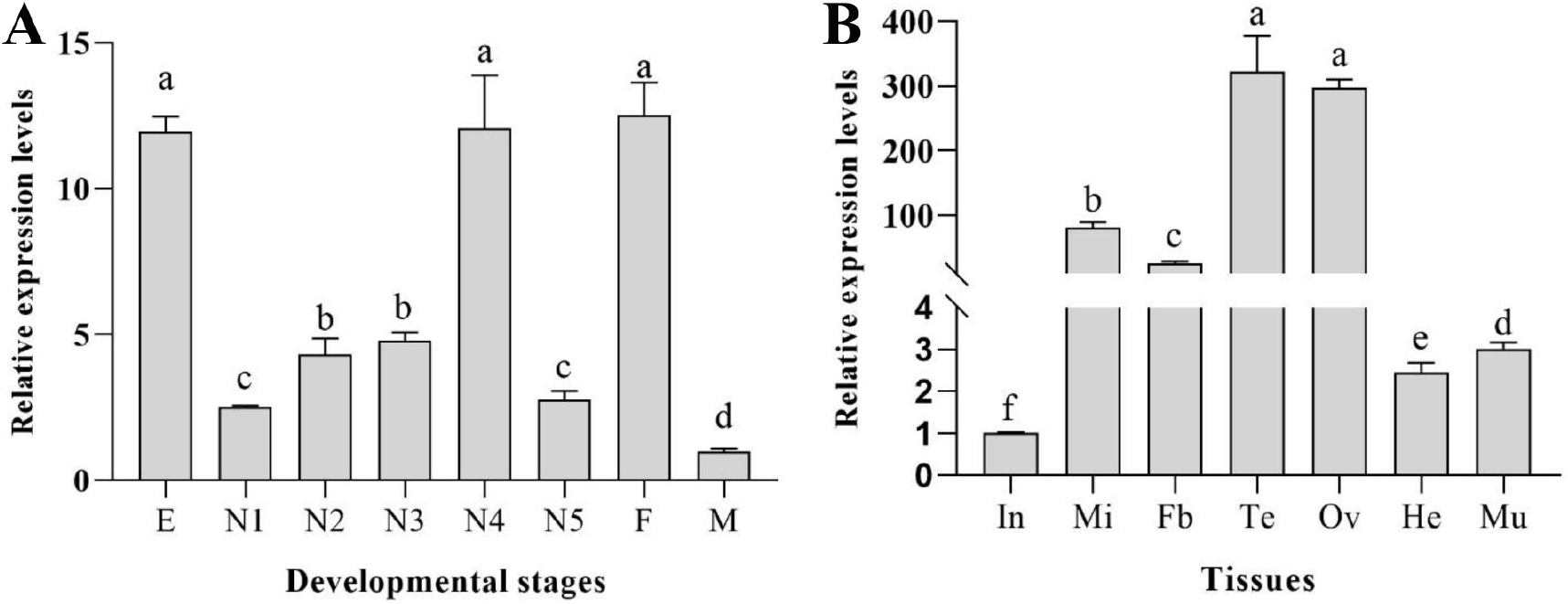
Relative expression levels of *CcAkt* at different developmental stages and in various tissues of *C. chinensis* adults. (A) Egg (E); 1st-5th instar nymphs (N1-N5); female (F); male (M). (B) Integument (In); midgut (Mi); fat body (FB); testis (Te); ovary (Ov); head (He); muscle (Mu). Values are the mean *±* SD of three replicates. Differences between groups were analyzed using one-way ANOVA and Duncan’s multiple range test. Different lowercase letters above the columns indicate significant differences (*p* < 0.05, Duncan’s test).

### 3.4. Effect of Exogenous Hormones on CcAkt, CcVg and CcVgR Expression

The expression profiles of *CcAkt, CcVg* and *CcVgR* after bovine insulin and 20-hydroxyecdysone (20E) injections were determined using RT-qPCR. *CcAkt* was significantly induced, with upregulation of 4.32, 1.68 and 2.21-fold at 3, 6 and 24 h after bovine insulin injection compared with the control, with no significant change at 12 h after injection (Figure 5A). In contrast, 20E significantly downregulated *CcAkt* expression (Figure 5B). Both exogenous hormones induced the *CcVg* expression. *CcVg* was significantly induced, with upregulation of 4.69, 22.66, 6.09 and 3.88-fold at 3, 6, 12 and 24 h after bovine insulin injection compared with the control (Figure 5C). *CcVg* was significantly induced, with up-regulation of 1.50, 33.21, 12.39 and 5.24-fold at 3, 6, 12 and 24 h after 20E injection (Figure 5D). Expression peaked at 6 h for both genes. *CcVgR* expression gradually increased from 3 to 24 h after injection, and peaked at 24 h. *CcVgR* was significantly induced, with upregulation of 23.49 and 5.28-fold at 24 h after injection of bovine insulin (Figure 5E) and 20E (Figure 5F), respectively, compared with the control.

**Figure 5.**
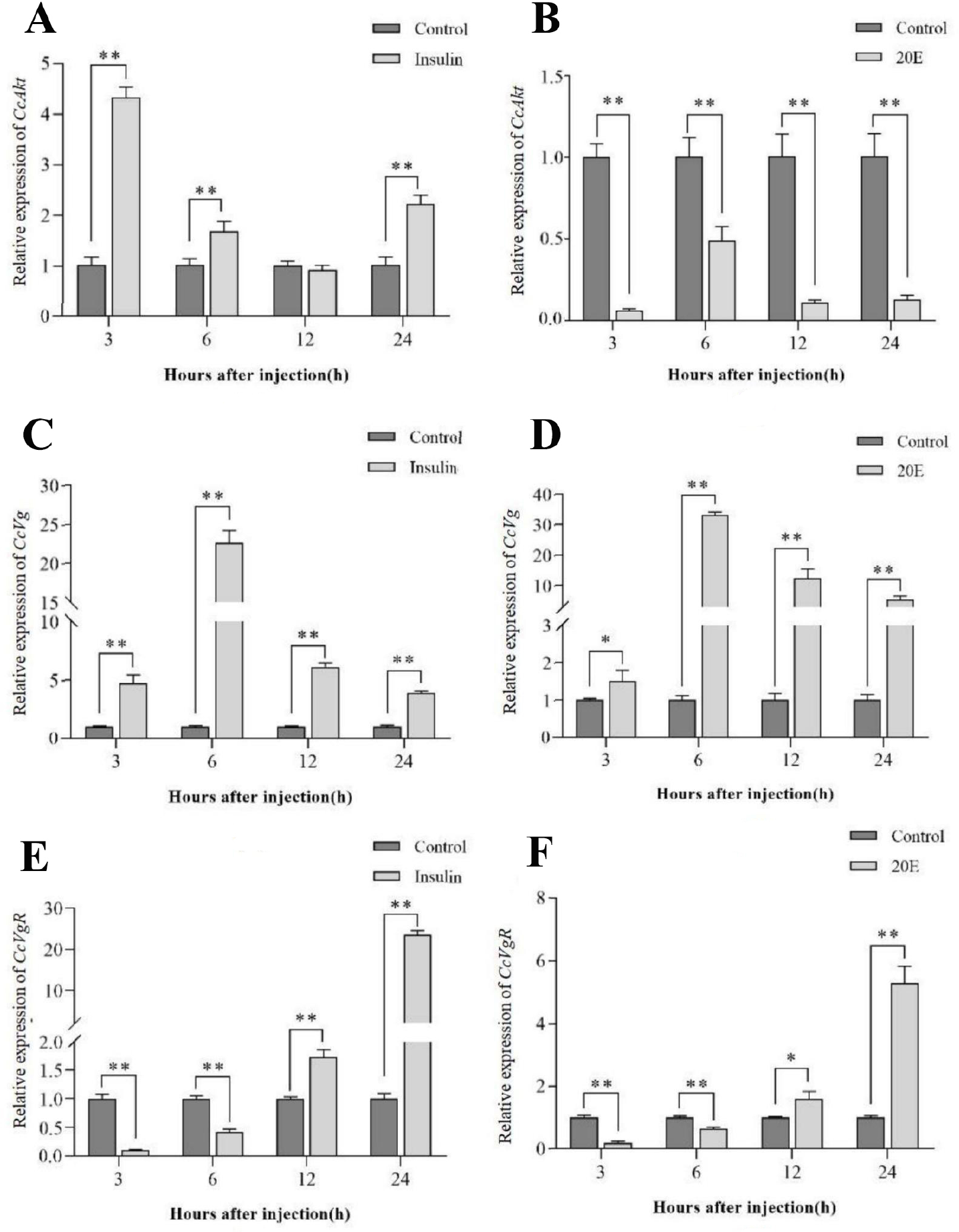
Expression profiles of *CcAkt, CcVg*, and *CcVgR* after exogenous hormone treatments. (A) Effect of bovine insulin on the expression of *CcAkt*. Control: insects injected with HEPES buffer. (B) Effect of 20E on the expression of *CcAkt*. Control: insects injected with distilled water containing 0.1% ethanol. (C) Effect of bovine insulin on the expression of *CcVg*. Control: insects injected with HEPES buffer. (D) Effect of 20E on the expression of *CcVg*. Control: insects injected with distilled water containing 0.1% ethanol. (E) Effect of bovine insulin on the expression of *CcVgR*. Control: insects injected with HEPES buffer. (F) Effect of 20E on the expression of *CcVgR*. Control: insects injected with distilled water containing 0.1% ethanol. Asterisks above the bars indicate significant differences between the treatment and the control (*, *p* < 0.05; **, *p* < 0.01).

### 3.5. Effect of dsCcAkt Injection on RNAi Efficiency

Using RT-qPCR, the assay results compared with the control group showed that *CcAkt* expression was reduced by 58.17%, 14.09%, 51.85% and 77.50% at 24, 48, 72 and 96 h after knockdown, respectively, demonstrating that *CcAkt* knockdown was a continuous process (Figure 6A). As the greatest decrease was observed at 96 h after gene silencing, samples were collected at 96 h after injection to examine the effect of silencing *CcAkt* on the expression of other insulin signaling pathway genes. *CcInR, CcFoxO, CcPDK, CcPI3K-C* and *CcPI3K-R* expressions were significantly reduced after silencing *CcAkt* compared with the control group, with downregulation of 61.20%, 84.86%, 67.84%, 76.46% and 33.90%, respectively. Overall, the expression of all insulin signaling pathway-related genes was suppressed after ds*CcAkt* injection (Figure 6B).

**Figure 6.**
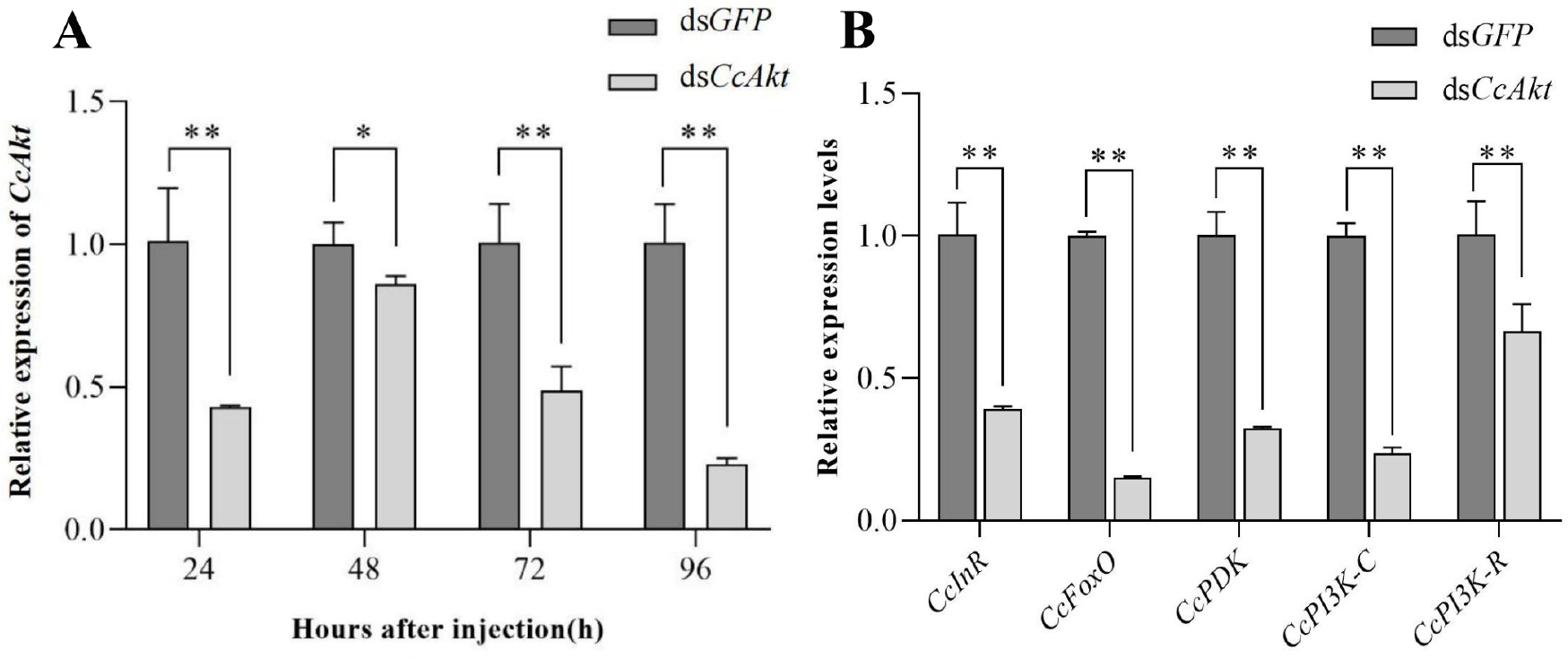
Effects of the *CcAkt* knockdown on expressions of the insulin signal pathway genes. (A) Relative expression levels of *CcAkt* after injection of dsRNA. (B) Relative expression levels of the insulin signal pathway genes 96 h after injection of ds*CcAkt*. Asterisks above the bars indicate significant differences between the treatment and the control (*, *p* < 0.05; **, *p* < 0.01).

### 3.6. Effect of Silencing CcAkt on Female Fertility and Ovarian Development

After demonstrating that the knockdown of *CcAkt* is a continuous process, we examined the *CcVg* and *CcVgR* expressions after the *CcAkt* knockdown to investigate the role of *CcAkt* in the reproduction of *C. chinensis. CcVg* and *CcVgR* expressions decreased by 37.93% and 78.04%, respectively, within 96 h after ds*CcAkt* injection compared with the control group (Figure 7A). On day 15 after ds*CcAkt* injection, ovaries were dissected from female adults and observed. *CcAkt* knockdown could stunt the development of the ovarioles compared with the control group (Figure 7B) and the length of ovarioles was significantly shorter (Figure 7C). As the knockdown of *CcAkt* resulted in incomplete ovarian development in female adults, we examined the parameters related to fertility after gene disturbance. Each ds*CcAkt*-treated female adult laid an average of 15.90 eggs compared with an average of 27.30 eggs in the control females injected with ds*GFP* (Figure 7D). The egg-hatching rate of ds*CcAkt*-injected females decreased by 9.91% compared to the control (Figure 7E).

**Figure 7.**
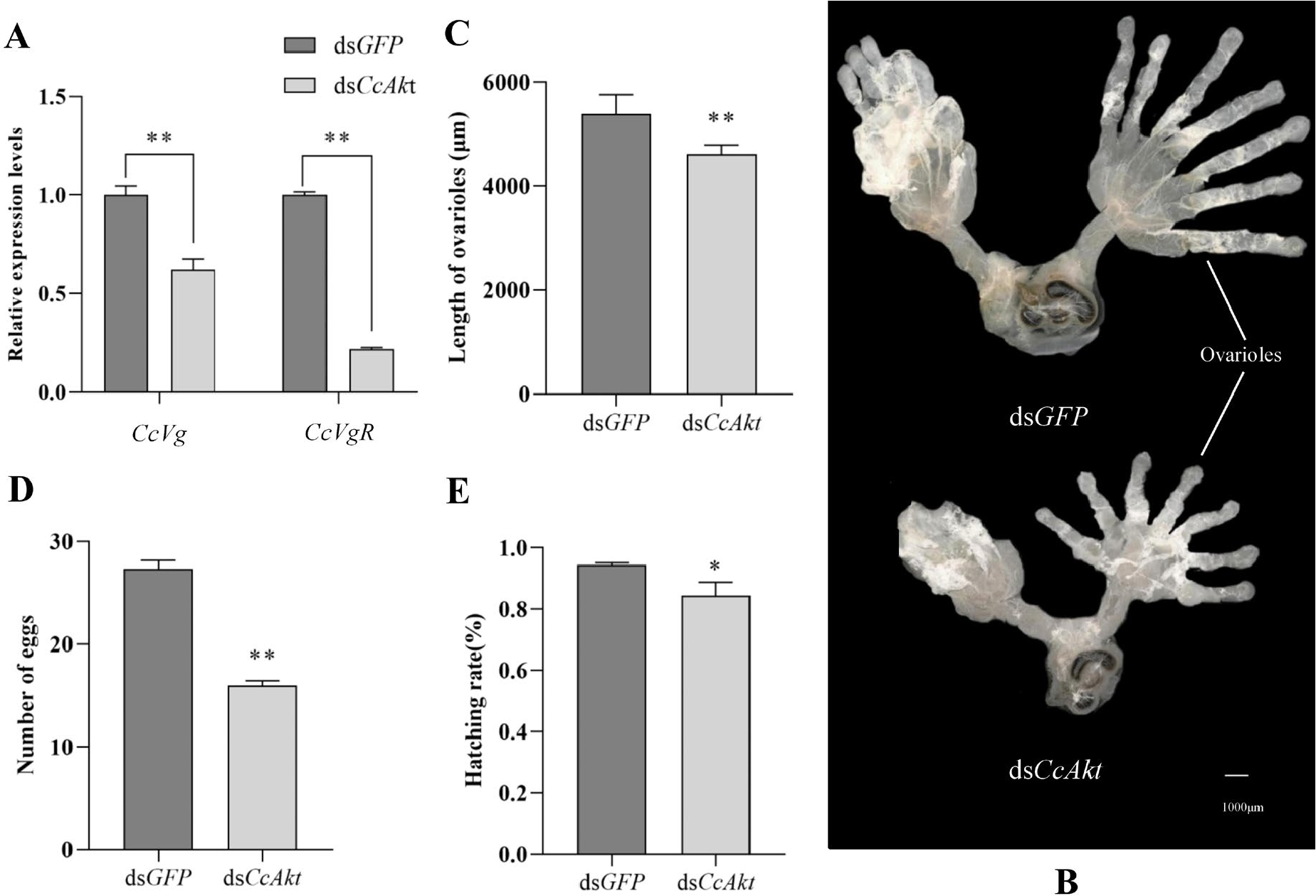
Effect of *CcAkt* by RNAi on fecundity of *C. chinensis*. (A) Effect of *CcAkt* by RNAi on the *CcVg* and *CcVgR* expressions. (B) Changes in ovarian morphology after injection of ds*CcAkt*. The ovaries of the ds*CcAkt*-injected females became smaller compared to the control group. Scale bar =1,000 μm. (C) Changes in length of the ovarioles after injection of ds*CcAkt*. The ovarioles in the ds*CcAkt*-injected females were much shorter than those in the control. (D) The number of eggs 15 d after injection of ds*CcAkt*. (E) The hatching rate of eggs after injection of ds*CcAkt*. Asterisks above the bars indicate significant differences between the treatment and the control (*, *p* < 0.05; **, *p* < 0.01).

## 4. Discussion

We examined the expression pattern of *CcAkt* at different developmental stages of *C. chinensis* and in different tissues of adult. The results showed that *CcAkt* was highly expressed in female adults and egg stages. The same researchers found in *A. aegypti* that *AaAkt* is expressed only in early embryonic and adult females [29]. We speculate that the insulin signaling pathway may participate in the embryonic development and reproductive regulation of *C. chinensis*. The expression level of *CcAkt* was higher at age 4 than at other ages, and it is speculated that the insulin signaling pathway may be involved in the regulation of insect wings. It has been shown that wing differentiation in *Nilaparvata lugens* is associated with the regulation of the insulin signaling pathway [30]. *CcAkt* expression also showed tissue specificity, with the highest expression in the testis and ovary compared to other tissues, confirming the crucial role of *CcAkt* in *C. chinensis* reproduction. In *A. aegypti, AaAkt* is most highly expressed in the ovaries of adult females [29]. This is generally consistent with our findings. However, the expression of *CcAkt* in other adult insect tissues should not be neglected, especially in the midgut and fat body. For insects, the midgut plays an important role in digesting food and absorbing nutrients as a major site for the secretion of digestive enzymes as well as other enzymes, while the fat body is an important site for nutrient storage and energy metabolism in the body, influencing insect cell and individual size through the regulation of hormonal and nutritional signals [31,32]. *DmAkt* was also detected in *D. melanogaster* with high expression in the fat body [33].

Insect reproductive development is a complex process that requires the involvement of multiple factors, including JH, 20E, insulin signaling, and other regulatory factors. Bovine insulin was found to directly stimulate the production of ecdysone in the ovary through the insulin signaling pathway, a finding reported in *A. aegypti* [34]. Similarly, in the prothorax of *B. mori*, it was found that injection of bovine insulin induced *BmAkt* expression and could lead to *BmAkt* phosphorylation, which in turn stimulated ecdysone secretion [11]. Bovine insulin treatment of *B. mori* larvae resulted in a significant increase in the expression of *BmInR, BmPI3K* and *BmAkt* [35], whereas 20E treatment of *Lasioderma serricorne* suppressed the transcriptional levels of *LsAkt* [36]. In *Chrysopa septempunctata*, injection of bovine insulin increased mRNA and protein levels of Vg, promoted ovarian growth, increased Vg protein abundance, improved reproductive performance, and enhanced protease activity in female insects [14]. In this study, bovine insulin injection promoted the expression of *CcAkt*, while 20E injection exerted an inhibitory effect on *CcAkt*. These two hormones upregulated the expression of *CcVg* and caused a gradual increase in the expression of *CcVgR*. These results suggest that both bovine insulin and 20E can participate in the reproductive regulation of insects by regulating the insulin signaling cascade response in insects.

In this study, *CcAkt* was silenced by RNAi and was detected to have different degrees of effects on five other important genes of the insulin signaling pathway, so we speculate that this may be related to the feedback and negative feedback regulatory mechanisms between genes of the insulin signaling pathway. Under special conditions, insects initiate specific stress mechanisms in vivo, generating a series of feedback and negative feedback regulatory mechanisms that will cause complementary effects between genes to maintain metabolic homeostasis in the insect [37]. Furthermore, silencing of *CcAkt* was found to have a suppressive effect on *CcVg* and *CcVgR* mRNA transcript levels, ovary size, egg production and hatching rate. In *D. melanogaster*, it was shown that the insulin signaling pathway directly regulates oocyte growth and that silencing key factors in the insulin signaling pathway prevents the uptake of yolk protein precursors and retards oocyte development [8]. In *M. vitrata*, interference with the insulin signaling pathway gene *MvAkt* was found to suppress the expression of *Vg* and *VgR* and impede normal ovarian development [9]. In *C. pallens*, RNAi to *CpAkt* significantly reduced *Vg* expression and led to ovarian atrophy and reduced yolk protein deposition [10]. These studies are consistent with the results of the present study, and we therefore infer that *CcAkt* is involved in the regulation of normal reproduction in *C. chinensis*.

## Funding

This work was supported by the National Natural Science Foundation of China (No. 31560610). The funder had no role in the study design, data collection and interpretation, or the decision to submit the work for publication

## Data Availability Statement

The datasets used and analyzed during the current study are available from the corresponding author upon reasonable request.

## Conflicts of Interest

The authors declare no conflict of interest.

## Notes

### Competing Interest Statement

The authors have declared no competing interest.

